# Carbon Stock of Human-Disturbed Forest Areas in Bukidnon, Philippines

**DOI:** 10.1101/2020.04.14.041798

**Authors:** Michael Arieh P. Medina, Vember C. Cabahug, Gabriel Erico G. Zapico

**Affiliations:** Department of Environmental Science, College of Forestry and Environmental Science, Central Mindanao University, University Town, Musuan, Bukidnon, Philippines

**Author notes:** Author to whom all correspondence should be addressed.

**Keywords:** carbon stock, biomass, forest, carbon sequestration

## Abstract

A carbon stock assessment was done on two forest areas in Bukidnon, Philippines specifically in Mt. Kiamo in Kibalabag Village and Mt. Capistrano in Managok Village both in Malaybalay City, Bukidnon. Using a nested sampling design, tree diameter at breast height as well as tree height were measured. Allometric equations were then used to calculate for the above and below ground biomass density of trees. Destructive sampling was then employed to determine the biomass density of understory and litterfall, while composite sampling was done for soil analysis. Carbon content was then used to compute for the carbon stock of the different carbon pools in the studied ecosystems. The average biomass density of the two areas was found to be 247.80 Mg ha^−1^ which are closely similar to other areas of the same forest type in Mindanao. Likewise the average carbon stock of the two areas is 143.14 MgC ha^−1^ which is similar to tree plantations based on previous studies. Furthermore, the said carbon stock is approximately 40% lower than the country average for natural forests (250 MgC ha^−1^). The results demonstrate the extent of the diminishing carbon sequestration capability of forest areas due to human disturbance.

## 1. Introduction

It has been reported that global warming is now undeniable, in fact since the 1950s; many of the observed changes in the climate system are unprecedented over decades to millennia. It is also a fact that these changes in the climate system brought about by global warming can be attributed to the increased concentrations of greenhouse gases in the atmosphere. Due mainly to human activities, the concentrations of greenhouse gases have exceeded the pre-industrial levels by about 40%, 150%, and 20%; specifically for carbon dioxide, methane, and nitrous oxide, respectively (IPCC, 2013).

Moreover, forestry related approaches such as afforestation, sustainable forest management, and reducing deforestation have been recognized as cost-effective climate change mitigation options though with large differences in their relative importance across regions (IPCC, 2014). Forests serve as a vast carbon sink thereby offsetting the carbon emission especially from human activities (Lasco et al, 2008). On an average, 160 Gt of carbon have accumulated in terrestrial ecosystems (including forests) globally, from 1750 to 2011 (IPCC, 2014).

The Philippines is blessed with vast forest areas which are estimated to be around 15.9M hectares. These forest areas leads to total land use change and forestry (LUCF) sector sequestration in the country which is almost equal to its total net GHG emission from all sources (101 Mt in 1994). This is an evidence of the importance of Philippine forests in climate change mitigation, because they absorb practically all the fossil fuel emissions of the country (Lasco and Pulhin, 2003). However, ongoing human disturbances in these forest areas threaten the carbon sequestration capacities of these ecosystems. It has been proven that forest disturbance is associated with the decrease in forest biomass density hence also decreasing carbon stock (Yohannes et al, 2015).

In line with the above facts, it is the aim of this study to determine carbon stocks among human-disturbed forest areas in the country specifically in its southern parts where forest degradation is rampant. Information that will be generated from this study hopefully will provide facts on the extent of the effect of human disturbances on forests specifically on its climate change mitigation capacity.

## 2. Materials and Methods

### 2.1. Study Location

Two forest areas were considered in the study. The first one is in Mt. Kiamo (8°17’26.3“N 125°09’15.0”E) and the second one is in Mt. Capistrano (8°00’35.9“N 125°10’45.6”E). Both are located in Malaybalay City, in the province of Bukidnon, Philippines (Figure 1). Bukidnon is an agricultural province which is part of the island of Mindanao, the third largest island group in the Philippines. The climate classification of Malaybalay City falls under Fourth Type or intermediate B type which is characterized by evenly distributed rainfall throughout the year, with no distinct dry season and maximum rain period observed is from October to February.

**Fig. 1.**
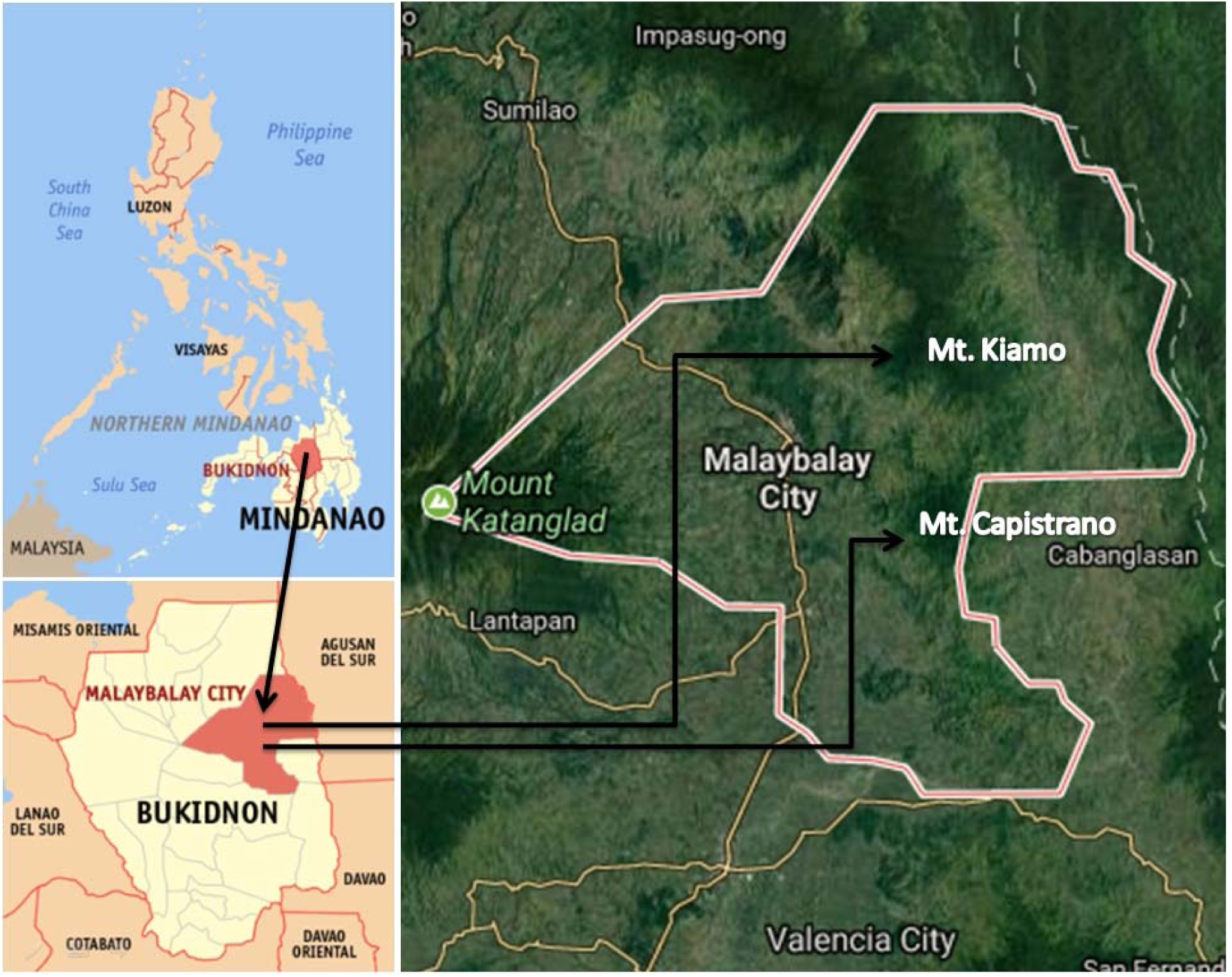
Location map of the study areas (Wikimedia Commons and Google Earth)

Mt. Kiamo has two distinct habitat types of ecosystem: the forest and grassland. Grassland area can be observed within the range of 1,193m to 1,400m above sea level while the forested areas can be observed within the range of 1400m – 1790m above sea level, and characterized as primary mature forest. Moreover, grassland areas are found along the trail towards the forest this is characterized as very steep dominated primarily by cogon and shrubs. On the other hand, Mt. Capistrano is a 610 meters high land form crowned with magnificent steep, sharp rocks. The trail going to the summit comprises of natural as well as secondary growth forests. The distance between Mt. Kiamo and Mt. Capistrano is roughly 30km.

Currently in Mt. Kiamo, several patches of forests are cleared for agriculture. Mostly, forest settlers are practicing slash and burn farming which is seen to greatly affect the essential forest resources of the mountains. On the other hand, Mt. Capistrano also suffers the same fate as evidenced by some cultivated lands within its forests. Furthermore, both areas are also seen as potential tourist spots especially to hikers and adventurists thus threatening further the ecological resources of the said areas.

### 2.2. Sampling Plot Establishment

A nested sampling design developed by Hairiah et al. (2001) was used in this study (Figure 2). Six and eight sampling plots each measuring 5mx40m were established in Mt. Kiamo and Mt. Capistrano, respectively. Field measurement were then taken in the different carbon pools using the procedures that follow.

**Fig. 2.**
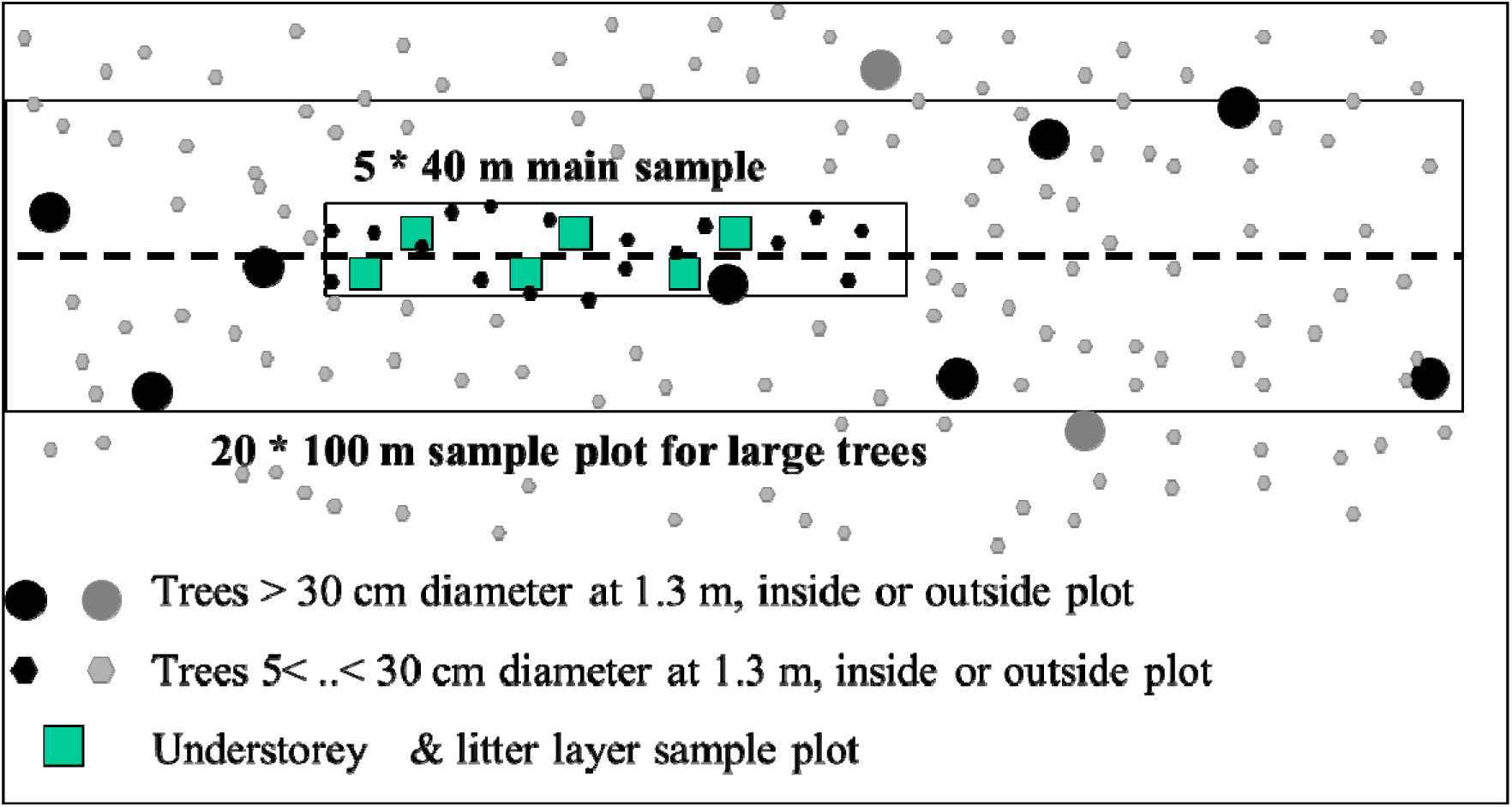
Field layout of nested sampling design (Hairiah et al., 2001)

### 2.3. Live Tree Biomass

The field sampling protocol was adopted from Hairiah et al. (2001). For live tree biomass measurements, a 40m transect line was laid through the center of the 5mx40m plot. All trees more than 5 cm diameter at breast height (DBH) within 2.5m of each side of the 40m transect line were measured using a tree caliper. For each tree, the diameter was measured at 1.3m above the soil surface except where trunk irregularities at the height occur and require measurement at a greater height. If trees with greater than 30cm in DBH were present in the sampling plot, whether or not they were included in the transect, an additional larger plot of 20mx100m (2,000m^2^) was established to include all trees with DBH greater than 30cm.

Aboveground biomass density of trees was calculated using the allometric equation below by Brown (1997).

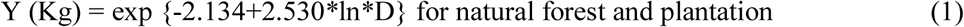

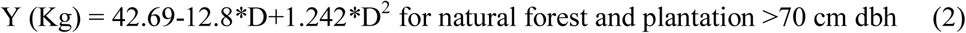

Where:

Y= above ground tree biomass, kg/tree

Exp= raise to the power of

In= natural logarithm

D= tree diameter at breast height (DBH), cm

Tree biomass density in tons per hectare is then computed using the equation below:

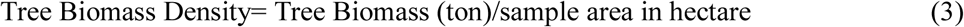

Consequently, carbon stored in the tree biomass was calculated by multiplying biomass density with the percent carbon content of trees. A default value of 0.45 (45%) was used for the carbon content of trees based on the average carbon content of wood samples collected from secondary forests from several locations in the Philippines (Lasco and Pulhin, 2000).

### 2.4. Understory Biomass and Litter layer

For understory and litter biomass, destructive sampling techniques were used. Within the 5mx40m plots, 1mx1m subplots were randomly placed in each quarter of the length of the 40m center transect line (Figure 2). Understory biomass which included trees <5cm DBH and all herbaceous vegetation, vines and lianas were harvested within the 1×1m subplots. The total fresh sample was weighed in the field after which a sub-sample of about 300g was taken for subsequent oven-drying and carbon content analysis. For litter layer the undecomposed plant materials or crop residues including all unburned leaves and branches were collected in 0.5mx0.5m quadrat on a random location within the understory sample plot (Figure 2). The undecomposed (green or brown) materials were weighed and then a sub-sample of about 300g was also taken for oven drying and carbon content analysis.

The ground samples that were taken during the data gathering were exposed to air and oven drying and grinding. Biomass of understory and litter were calculated using the following formula:

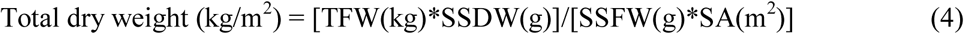

Where:

TFW = Total fresh weight

SSDW = Sub-sample dry weight

SSFW = Sub-sample fresh weight

SA = Sample area

Furthermore, carbon stored in the understory and litter biomass was calculated by multiplying total dry weight with the percent carbon content of samples from both carbon pools. Carbon content values were based on the results of the carbon content analysis.

### 2.5. Root Biomass

Since the root biomass of trees are difficult to estimate as it involved destruction, the following allometric equation was used based on aboveground biomass for tropical forest [Cairns et al., 1997).

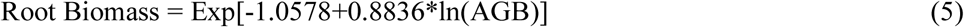

Where:

Exp= e to the power of

In= Natural logarithm

AGB= Aboveground biomass

### 2.6. Soil organic carbon

For soil organic carbon, the same sampling quadrats for litter sampling was used (Figure 2). A composite sample of about 500g of soil samples were taken from each 5×40 m plot. The samples were obtained at 0-30cm depth and were air dried. The air dried samples were then subjected to laboratory analysis to determine the soil organic carbon (SOC) using the Walkey-Black Method. Bulk density was determined by collection of undisturbed soil cores with a sampling metal tube with a diameter of 5.3cm and a length of 10cm. Said Samples were subjected for oven drying to constant weight for 40 hours at ± 105°C. Bulk density (BD) and soil organic carbon (SOC) was computed as follows:

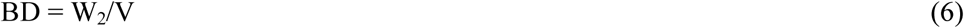

Where:

BD = Bulk density of the soil sample

W_2_ = Oven dry weight of the sample

V = Volume of the cylinder/tube

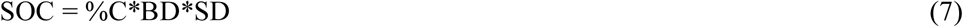

Where:

SOC = Soil Organic Carbon (MgC/ha)

BD = Bulk Density (Mg/m^3^)

SD = Sampling Depth (cm)

## 3. Results and discussion

### 3.1. Biomass Density of the Study Areas

As shown in Table 1, biomass density of forest in Mt. Kiamo and Mt. Capistrano equates to 195.55 Mg ha^−1^ and 300.06 Mg ha^−1^, respectively. With this, it is evident that Mt. Capistrano is considered to have a higher biomass density compared to Mt. Kiamo. Spatial differences in biomass of forests have been explained to be caused by factors such as soil, microclimate, structure, and human disturbance (Clark and Clark, 2000; Ensslin et al., 2015; Paoli et al., 2008; Sheikh et al., 2009). In the case of the study, Mt. Kiamo is observed to show evidences of human disturbance as it is currently threatened by ongoing slash and burn agriculture in several of its patches of forest. Mt. Capistrano, on the other hand has lesser threats in comparison in the context of the above human disturbances, however, recent tourism activities in the mountain such as trail hiking is seen to presently threaten the forest resource of the area in the near future unless it is managed properly.

**Table 1.**
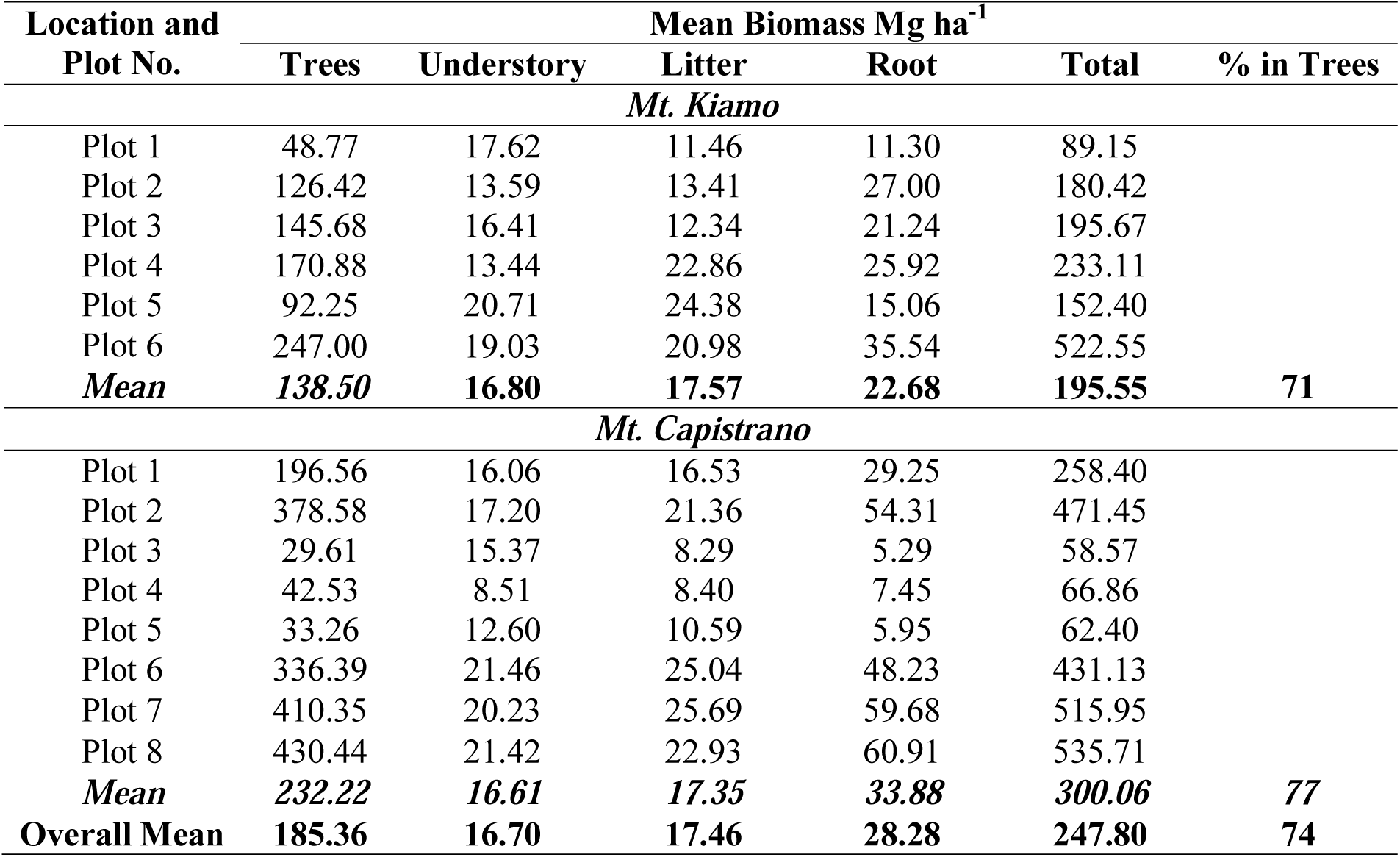
Biomass density (Mg ha-1) of different carbon pools

The mean biomass density of both forest areas in the study is 247.80 Mg ha-^1^. This value is a closely similar to the biomass density of second growth forests in Mindanao (262.0 Mg ha^−1^) as reported by Lasco and Pulhin (2003). However, in the same report, second growth forests in Mt. Makiling in Luzon have twice as much biomass density (547.0 - 672.8 Mg ha^−1^). On the overall however, based on the same report, most forest tree plantations have lower biomass density than the ones in this study.

Furthermore, majority of the biomass density of the study areas is found in trees (74%). This actually supports the results in previous studies of plantations in Bukidnon (Janiola and Marin, 2016; Labata et al., 2012; Patricio and Tulod, 2010; Tulod, 2015) in which trees hold as low as 58% of the total biomass in tree plantations to as high as 94% in agroforestry farms. Tree trunks are not only larger compared in size to the other biomass components but also they live long thus they can store the most carbon in a particular forest (Patricio and Tulod, 2010). On the other hand, roots are the second highest biomass storage in the study followed by litter then understory. Same trend is observed with values in the different biomass components from the previously mentioned studies (Janiola and Marin, 2016; Labata et al., 2012; Patricio and Tulod, 2010; Toledo-Bruno et al., 2017; Tulod, 2015).

### 3.2. Carbon Stock in the Study Areas

As shown in Table 2, Mt. Kiamo has an average carbon stock of 128.97 MgC ha^−1^ while Mt. Capistrano has an average carbon stock of 157.31 MgC ha^−1^. As observed, carbon stock values follow the same variation as biomass, meaning the higher the biomass, the higher the carbon stock. Furthermore, the average carbon stock for both areas is 143.14 MgC ha^−1^. The findings shows that the carbon stock in both areas are comparable to forest tree plantations (Patricio and Tulod, 2010; Tulod, 2015), as well as agroforestry farms (Labata et al., 2012), and fruit tree plantations (Janiola and Marin, 2016) in areas located within the province of Bukidnon (Table 3). On a national level, the same carbon stock values also fall within the previously observed carbon stock values in forest plantations (Lasco and Pulhin, 2003) yet it is 40% lower than the average for natural forests (250 MgC ha^−1^) in the country (Lasco et al., 2008). This just proves the threatened condition of these areas.

**Table 2.** Carbon stocks (MgC ha^−1^) in the different carbon pools.

| Location | Mean Carbon Stocks (MgC ha<sup>-1</sup>) |  |  |  |  |  |
| --- | --- | --- | --- | --- | --- | --- |
|  | Trees | Understory | Litter | Root | Soil | Total |
| Mt. Kiamo | 64.83 | 8.56 | 9.30 | 10.21 | 36.07 | 128.97 |
| Mt. Capistrano | 104.50 | 8.12 | 7.72 | 15.25 | 21.72 | 157.31 |
| **Mean** | **84.66** | **8.34** | **8.51** | **12.73** | **28.90** | **143.14** |

**Table 3.** Carbon stocks of different plantation areas in Bukidnon, Philippines

| Location | Forest/Plantation Type | Carbon Stock (MgC ha<sup>-1</sup>) | Reference |
| --- | --- | --- | --- |
| Malaybalay and Impasug-ong | Benguet Pine Plantation | 158.98 | Tulod and Patricio, 2010 |
| Musuan, Maramag | Teak Plantation | 226.80 | Tulod, 2015 |
| Musuan, Maramag | Second Growth Forest | 217.46 | Tulod, 2015 |
| Musuan, Maramag | Mahogany Plantation | 194.42 | Tulod, 2015 |
| Musuan, Maramag | Mangium Plantation | 163.22 | Tulod, 2015 |
| Musuan, Maramag | Yemane Plantation | 112.72 | Tulod, 2015 |
| Songco, Lantapan | Mixed Multistorey System | 161.52 | Labata et al, 2012 |
| Musuan, Maramag | Taungya Agroforestry System | 174.43 | Labata et al, 2012 |
| Panalsalan, Maramag | Falcata-Cacao Multistorey | 92.33 | Labata et al, 2012 |
| Musuan, Maramag | Rambutan Plantation | 112.18 | Janiola and Marin, 2016 |
| Musuan, Maramag | Mango Plantation | 122.34 | Janiola and Marin, 2016 |
| Musuan, Maramag | Santol Plantation | 203.62 | Janiola and Marin, 2016 |
| Kibalabag, Malaybalay | Disturbed Natural Forest | 128.97 | This Study |
| Managok, Malaybalay | Disturbed Natural Forest | 157.31 | This Study |

The above results suggest the substantial potential of the study areas for climate change mitigation through carbon storage and sequestration. However, the undeniable vulnerability of the above areas especially from human disturbance eventually downgrades its climate mitigation capacity.

Soil is observed to be the second major carbon pool (next to trees) which follows the same trend as the previous studies mentioned. Soil is an essential carbon pool since it is not released by burning and at the same time it has the longest residence time among the different carbon pools (Lugo and Brown, 1992). Furthermore, mean soil organic carbon in the study is 28.90 MgC ha^−1^. This result shows that carbon in forest soils as in the case of this study are higher than in agricultural soils (Patricio, 2014). Compared to agricultural lands, forests have higher organic matter inputs from roots and litters leading to higher soil organic carbon (Brown et al., 1991). As observed in the study Mt. Kiamo have comparably higher litter carbon (9.30 MgC ha^−^1) compared to Mt. Capistrano (7.72 MgC ha^−1^) which also explains the higher soil organic carbon in the former (36.07 MgC ha^−1^) compared to the latter (21.72 MgC ha^−1^). This implies that forest conversion into agricultural lands greatly contributes to climate change not only due to carbon released from the cutting of trees but also due to the degradation of carbon in soil.

## 4. Conclusions

The average biomass density for the threatened natural forests in the study is 247.80 Mg ha^−1^. Majority of the biomass are found in trees followed by roots, litter, and then understory. Furthermore, the average carbon stock in the study areas is 143.14 MgC ha^−1^.This is similar to carbon stored in several man-made forests within the province. However, it is found to be 40% lower than its expected carbon stock based on the average for natural forests in the country. This suggests the substantial capacity of such areas for climate change mitigation however, its vulnerability to human disturbance leads to its diminished potential for carbon sequestration.

It is hoped that this results will provide insights in encouraging the local government as well as private entities in the protection and conservation of Mt. Kiamo and Mt. Capistrano. Aside from the obvious ecological services provided by both areas (i.e. biodiversity, water provision, etc.), climate change mitigation through carbon storage and sequestration is a timely thus effective motivating factor in soliciting public support especially in the crafting of programs and policies related to the proper management of the said areas.

## Acknowledgement

The researchers wish to thank the village chiefs of Kibalabag (Mt. Kiamo) and Managok (Mt. Capistrano) in Malaybalay City, Bukidnon for graciously allowing the conduct of the study in their areas of jurisdiction. Dr. Jose Hermis P. Patricio and Dr. Angela Grace Toledo-Bruno of the CMU-CFES Department of Environmental Science have been instrumental in the completion of this research paper.

## Notes

### Competing Interest Statement

The authors have declared no competing interest.

